# Targeted 3’-end RNA sequencing uncovers cryptic polyadenylation in Huntington’s disease linked to somatic instability and CAG repeat purity

**DOI:** 10.64898/2026.01.03.697463

**Authors:** Ane Velasco-Bilbao, María Manterola, Alvaro Herrero-Reiriz, María Carazo-Hidalgo, Anna Misiukiewicz, Olatz Arnold-Garcia, Esther Perez-Navarro, Martina Hallegger, Jernej Ule, Alberto Rabano, Adolfo López de Munain, Marta Olejniczak, Veronica Brito, Lorea Blazquez

## Abstract

Huntington’s disease (HD) is a progressive neurodegenerative disorder caused by expanded CAG repeats in the first exon of the *HTT* gene, which encodes for huntingtin (HTT) protein. Full-penetrance is established at 40 repeats, but beyond, somatic repeat instability in the brain and CAG repeat purity modulate disease onset and severity. Previous studies have described that expanded repeats induce the incomplete splicing of *HTT* intron 1 to express the most pathogenic HTT isoform, known as HTT1a. Yet, the lack of a robust and sensitive method to evaluate *HTT* RNA-misprocessing has limited our understanding of *HTT1a* expression in HD pathophysiology.

Here we describe a targeted RNA sequencing approach, known as 3’-end targeted RNA sequencing or 3TRS, to simultaneously quantify multiple *HTT* transcripts generated by canonical and cryptic polyadenylation in several HD models. We show that activation of *HTT* cryptic polyadenylation is highly selective and requires long and uninterrupted CAG repeat expansions. In HD knock-in mice and human postmortem brain, cryptic *HTT* expression strongly correlates with brain-specific somatic repeat instability, supporting a model where ultralong and unstable CAG repeats drive toxicity by activating *HTT* RNA-misprocessing. Overall, 3TRS provides a robust framework to investigate *HTT1a* biogenesis and expression and to evaluate *HTT*-lowering therapeutic strategies.

**GRAPHICAL ABSTRACT:** 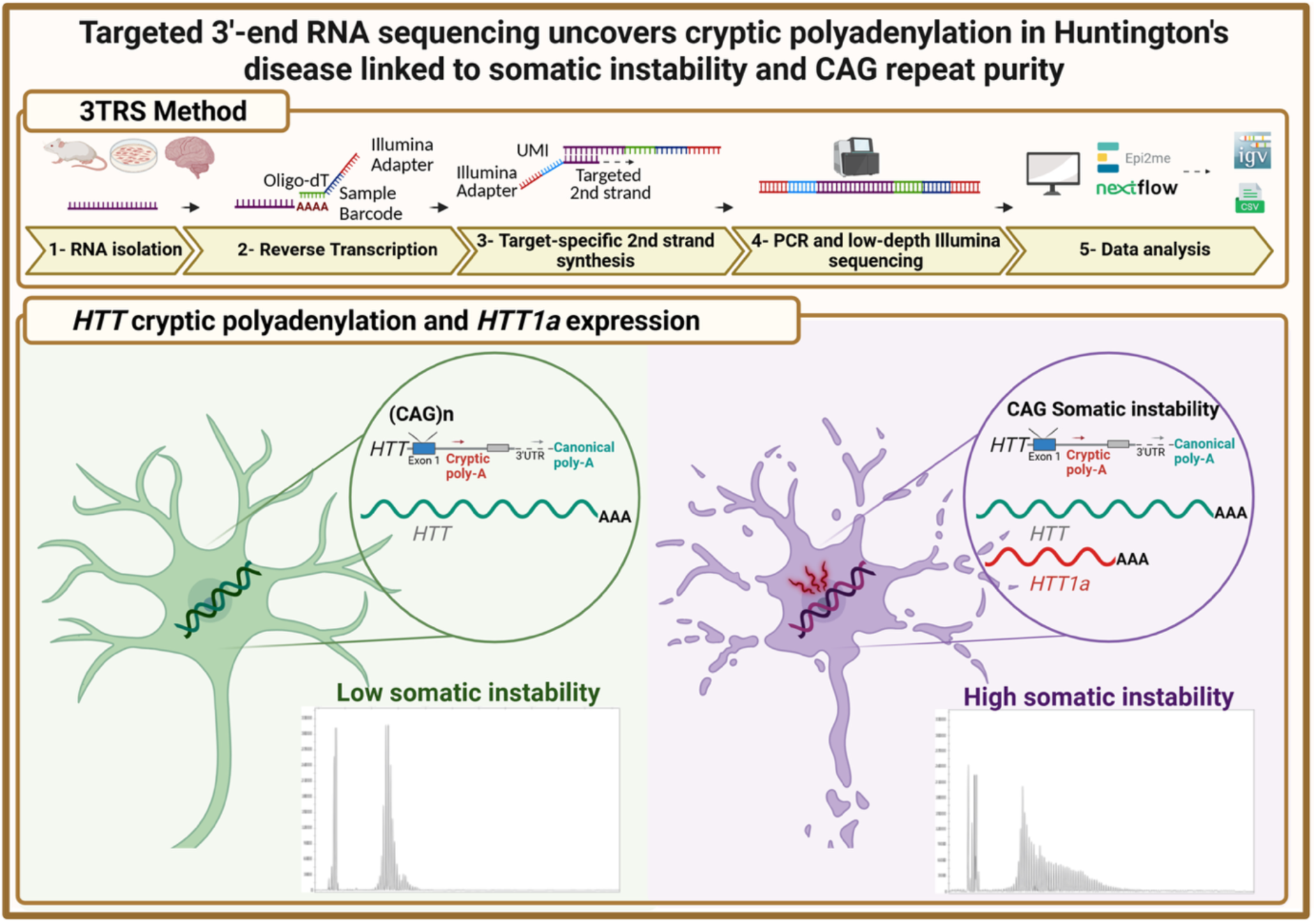

## INTRODUCTION

HD is a progressive neurodegenerative disorder with an autosomal dominant inheritance pattern caused by a CAG trinucleotide expansion in the first exon of the *HTT* gene (1). The repeat threshold to develop HD with full penetrance is established at ≥40 CAG repeats, although recent studies have demonstrated that ultralong somatic repeat expansions are required to drive toxicity in the brain (2–6). In *HTT*, the CAG repeat expansion is transcribed and translated to a polyQ tract within the huntingtin protein (HTT). HTT can be processed into multiple N-terminal fragments (7). Amongst them, the exon 1 HTT fragment known as HTT1a is considered the most pathogenic, and its expression is enough to produce the most aggressive HD mouse models (8, 9). Although several proteases that generate HTT N-terminal fragments have been identified, the enzyme responsible for the biogenesis of the smallest and most toxic exon 1 HTT fragment is still undetermined (10–13).

RNA misprocessing is a molecular hallmark of human disease, which involves alterations in RNA splicing and 3’-end processing (14, 15). While RNA mis-splicing is a well-established feature of neurodegeneration (16), the implication of aberrant 3’-end processing in disease has been investigated in less detail. In the context of HD, previous work from the Bates group has shown that expanded CAG repeats above a pathologic threshold lead to incomplete splicing of *HTT* intron 1 to generate an exon 1-intron 1 mRNA transcript. This isoform, known as *HTT1a*, is translated into the highly pathogenic exon 1 HTT fragment (17–19). Using several mouse and human HD models, several cryptic polyadenylation sites (pA-site) were described by rapid amplification of cDNA ends (3’RACE). Moreover, RT-PCR, quantitative PCR (qPCR), digital PCR and probe-based assays have been developed to measure *HTT* intron 1 fragments (17–22). Although informative, these assays do not rely on RNA sequencing, which limits the interpretation of the results, especially when low copy numbers of aberrant transcripts are generated (19, 20).

HTT lowering strategies are a major focus of therapeutic development for HD (23), especially those targeting *HTT1a* isoform with antisense RNA molecules (24, 25). To develop successful therapies, it is crucial to apply quantitative methods that accurately measure the expression of the targeted transcript, also in the context of all *HTT* isoforms. In this work, we performed targeted 3’-end RNA sequencing to measure all *HTT* transcripts in a sensitive, quantitative and cost-efficient manner. With this approach, we confirmed *HTT/Htt* cryptic polyadenylation in several human and mouse HD models. In humanized mouse models and cell lines, we clearly demonstrate that the distal cryptic pA-site, located 7.3kb within intron 1, is the one activated in the presence of an expanded CAG tract. In mouse *Htt*, however, both cryptic pA-sites within *Htt* intron 1 are used to a similar extent, specially upon somatic repeat expansion in the brain. Our data also demonstrate that CAA interruptions within the repeats block the expression of HTT1a transcript. In human HD samples, *HTT* cryptic polyadenylation occurs only in the presence of very long CAG tracts in a cell line from a juvenile-onset HD patient, or upon somatic instability in specific brain regions of postmortem tissue. Our data support a model in which very long, uninterrupted CAG-repeat expansions drive HD pathogenicity activating *HTT* intron 1 cryptic polyadenylation and HTT1a expression.

## MATERIAL AND METHODS

### Ethics approval and consent to participate

All experiments in this study were performed in accordance with the relevant guidelines and regulations. All mouse procedures were performed in compliance with the National Institutes of Health Guide for the Care and Use of Laboratory Animals and approved by the local animal care committee of the Universitat de Barcelona (448/17) and the Generalitat de Catalunya (9878 P2 and 9386), in accordance with the European (2010/63/EU) and Spanish (RD53/2013) guidelines for the care and use of laboratory animals. Procedures with fibroblasts from human origin were approved by the Ethics Committees of the Hospital de la Santa Creu I San Pau de Barcelona and the Universitat de Barcelona (IRB00003099, 07/20/2023), and informed written consent was obtained from all subjects. Human postmortem brain tissue from HD patients (Sup. Table 1) was obtained from Fundación CIEN, after the approval of the Clinical Research Ethics Committee of the Basque Country (code LBG-NDD-2023-01/23-30). Informed consent was obtained from all subjects involved in the study.

### Animals

Wild-type (WT), YAC128 (26) and heterozygous *Hdh^Q111/+^* knock-in (KI) mice (27) were kindly obtained from Dr. Verónica Brito (UB, Universidad de Barcelona). Mice were maintained on a C57BL/6J genetic background and housed with access to food and water ad libitum in a colony room kept at 19–22 °C and 40–60% humidity under a 12:12 h light/dark cycle. KI mice present a targeted insertion of 109 CAGs in the murine huntingtin gene that extends the resulting polyglutamine segment to 111 residues. The CAG repeat size for the KI mice used in this study was 112–119. To obtain age matched WT and *Hdh^+/Q111^* littermates, male WT mice were crossed with female heterozygous *Hdh^+/Q111^* mice. Only males from each genotype were used for the experimental procedures. Animals were sacrificed at 7 months of age, and brains were rapidly frozen in dry ice and stored at −80 °C until further analysis. All mouse procedures were performed in compliance with the National Institutes of Health Guide for the Care and Use of Laboratory Animals and approved by the local animal care committee of the Universitat de Barcelona (448/17) and the Generalitat de Catalunya (9878 P2), in accordance with the European (2010/63/EU) and Spanish (RD53/2013) guidelines for the care and use of laboratory animals.

### Cell lines and RNA extraction

Engineered HEK293T cells, homozygous with different numbers of CAG repeats at the *HTT* locus, were previously generated and published (21) while the HEK293 lines containing CAA interruptions were generated from a HEK293T wild-type cell with CRISPR-Cas9 system and contain a 41 poly-Q tract in homozygosis with one or two CAA interruptions across the CAG expansion. Both cell lines were kindly provided by Dr. Marta Olejniczak (Polish Academy of Sciences, Poland) and are listed in Sup. Table 2. Cells were cultured in Dulbecco’s modified Eagle’s medium (Gibco) supplemented with 10% fetal bovine serum (Teknovas) and 1% penicillin-streptomycin (Gibco) at 37 °C with 5% CO2. Cells were split when they reached 80% confluence and tested for mycoplasma using Venor PCR-based mycoplasma detection kit (MP0025). RNA from cultured HEK293T cells and fibroblasts was isolated using the Maxwell® RSC SimplyRNA Cells Kit (Promega, AS1390), according to the manufacturer’s protocol. RNA from mouse brain tissue was extracted using the RNeasy Lipid Tissue Mini Kit (Qiagen, 74804), as previously described (28). RNA from human brain samples was isolated using the Maxwell® RSC SimplyRNA Tissue (Promega, AS1340) or TRIzol™ reagent (Invitrogen, 5596026) after tissue homogenization with a pellet pestle (Sigma Aldrich, Z359963) or the OMNI Bead-Ruptor instrument. After RNA extraction, DNase treatment and RNA concentration was performed using the RNA Clean & Concentrator Kit (Zymo Research, R1013) following manufacturer’s instructions in a 15 µL final volume.

### *HTT* CAG repeat fragment analysis

100 ng RNA was converted to cDNA using the High-Capacity cDNA Reverse Transcription Kit (Applied Biosystems, 4368814) following manufacturer’s instructions. CAG repeat sizes were determined using a human-specific PCR assay (29), using the 6-FAM fluorescently labeled forward primer (5’-6FAM-ATGAAGGCCTTCGAGTCCCTCAAGTCCTTC-3’) (IDT), and reverse primer (5’-GGCGGCTGAGGAAGCTGAGGA-3’). 50 µL reactions were performed, using KOD Xtreme Hot Start DNA Polymerase (Sigma, 71975-M) according to manufacturer’s instructions. Cycling conditions were 2 min at 94 °C, 32 cycles of 10 seconds (s) at 98 °C, 30 s at 64 °C and 2 min at 68 °C. The reaction was then concentrated and purified using Nucleospin Gel and PCR clean-up (Macherey-Nagel, 740609), eluting the samples at 15 µL. Products were resolved using the SeqStudio Genetic Analyzer (Applied Biosystems), and GeneScan 1200 LIZ (Applied Biosystems, 4379950) was used as an internal size standard. CAG repeat size distribution plots were generated using GeneMapper v6 software. Somatic CAG expansion indexes were calculated as previously described (30), using a 10% relative peak height threshold cut-off. PCR products containing CAG repeats were sequenced by Plasmidsaurus (Germany, Europe), using the long-read amplicon sequencing service with Oxford Nanopore Sequencing Technology.

### Library preparation for 3’-end targeted RNA sequencing (3TRS)

For reverse transcription (RT), 500-1000 ng of each RNA (500ng for YAC128, 750ng for KI and 1 or 4ug for human samples) was used as starting material in a total volume of 11 µL. For each sample, 1 µL of reverse transcription primer at 250nM containing a sample barcode (Sup. Table 3) and 1µL of dNTPs at 10 mM (NEB, N0447L) were added. After 5 minutes of denaturation at 65 °C, samples were cooled to 37 °C and held in a thermocycler. Each sample was reverse transcribed using SuperScript IV reverse transcriptase. A mastermix containing 4 µL of SuperScript IV Buffer, 1 µL of 0.1M DTT, 1 µL of RNAsin® (Promega, N2615) and 1 µL of SuperScript IV reverse transcriptase (Invitrogen 18090050) was prepared and pre-warmed at 42 °C for 2-3 min before being added (7 µL / sample) to the samples held at 37 °C in the thermocycler. The RT reaction was then performed at 50 °C for 10 min and the reverse transcriptase enzyme was inactivated at 80 °C for 10 min and cooled to 25 °C. After this step, pools containing the RT reaction of six samples were mixed and purified. For purification, 2.5X Purification Beads (Lexogen SKU 022.96) were added, well mixed and incubated for 4-5min. Then, they were allowed to bind to the magnet (for 2-3min until the supernatant is clear), and the supernatant was taken off. The wash step was performed three times with freshly prepared 80% ethanol (500 µL) without taking the tube off the magnet. In the last wash, all the ethanol was removed (adding a spin step) and air-dried for 5 min. Finally, the sample was eluted by adding 50 µL of molecular biology grade water, incubated for 4-5 min, and allowed to bind for 4-5 min in the magnet. The sample containing the single-stranded cDNA (ss-cDNA) was collected in a new tube. For the second strand synthesis, 15 µL of High-Fidelity Phusion 2X Master Mix (NEB, M0531L); 13 µL of the ss-cDNA and 2 µL of the second strand synthesis oligos specific for our targets of interest (Sup. Table 3) were mixed in a final volume of 30 µL. The initial concentration of the second strand oligo mix was 7.5 µM independently of the number of targets multiplexed in each reaction. Each primer contains a unique molecular identifier (UMI), consisting of a random nucleotide sequence that allows demultiplexing PCR amplified cDNA reads to unique cDNA molecules. Initially, 6-nt long UMIs were employed for the data generated in Figures 1-3. Subsequent experiments were performed with 10-nt long UMIs to increase sequence complexity. The second strand synthesis reaction consists of a 2 min incubation at 98 °C; 60 s at 55 °C (expected annealing temperature for the primers) and a 5 min incubation at 72 °C to complete the second strand synthesis. At this point, samples were held at 10 °C and purified using 1X purification beads (Lexogen, SKU 022.96), following the manufacturer’s instructions. After the second strand synthesis, the double-stranded cDNA (ds-cDNA) was amplified by PCR. A common forward primer containing the Illumina sequencing adapter sequence was used for all the reactions. The reverse primer contained the Illumina sequencing adapter and a 6-nt-long index (i7) that differs for each oligo and allows indexed sequencing of each library (Sup. Table 3). The PCR reaction was set to a 30 µL final volume, containing 15 µL of the Phusion High-Fidelity 2X Master Mix (NEB, M0531L), 1.5 µL of each primer at 10 µM and 12 µL of purified ds-cDNA. The PCR program started at 98 °C for 30 s, followed by 25 cycles of 10 s at 98 °C, 30 s at 60 °C and 30 s at 72 °C. The size of the PCR product was first assessed by automated electrophoresis with the QIAxcel Advanced System using the QIAxcel DNA Screening kit (QIAGEN). Then it was purified from a 4% E-Gel^TM^ EX Agarose Gel (Invitrogen, G401004) using the E-Gel Power Snap Electrophoresis System (Invitrogen, G8300). In these gels, fragments between 150 bp and 400 bp were selected for band excision. The library was then purified using MiniElute PCR purification columns (Qiagen, 28004). Before sequencing, library size and integrity was analysed in the Agilent TapeStation using the High Sensitivity D1000 ScreenTape Assay (Agilent, 5067-5584) and library concentration was quantified with the Qubit™ dsDNA High Sensitivity Assay Kit (Invitrogen, Q32851).

**Figure 1:**
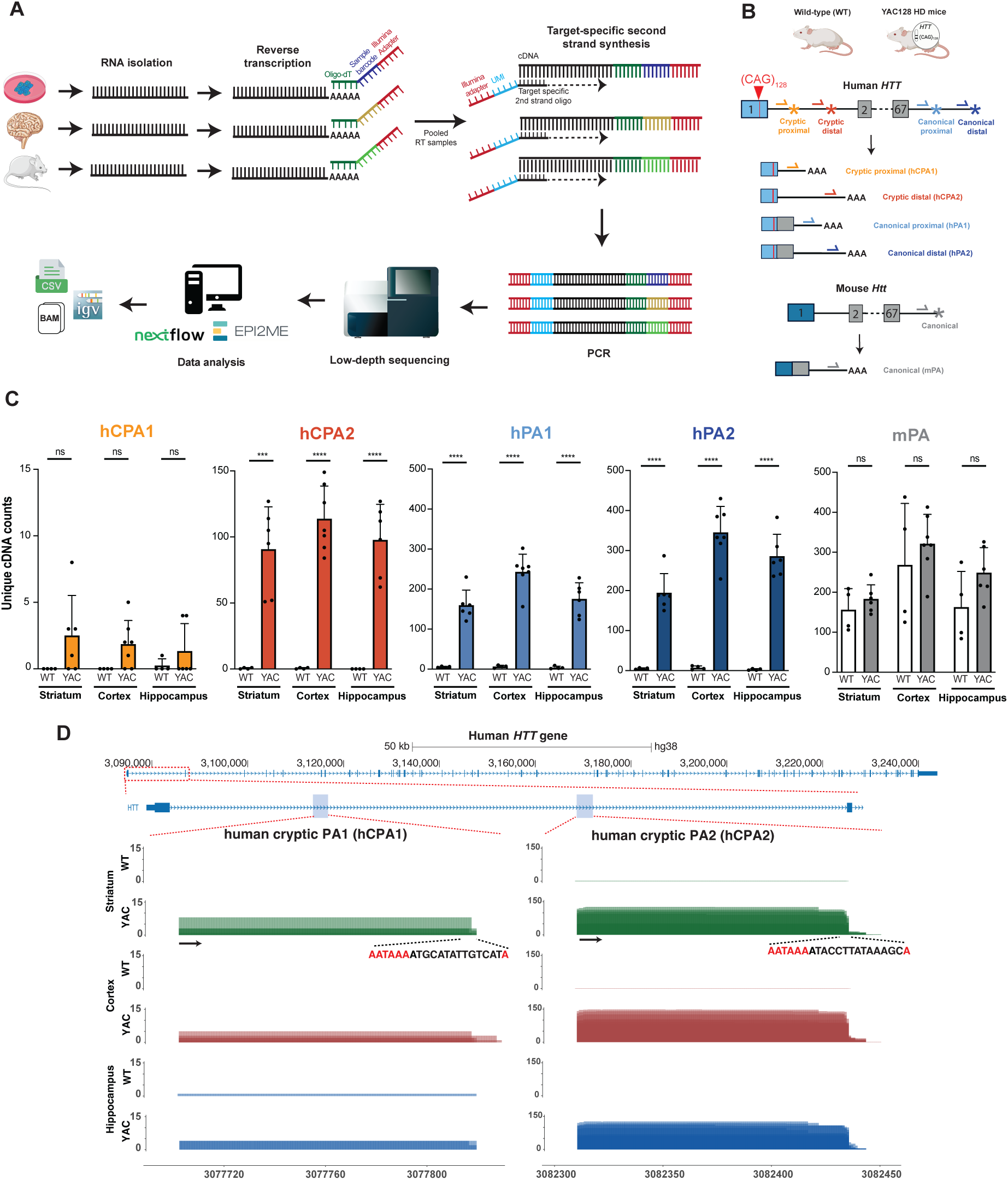
*HTT* RNA processing in the YAC128 mouse model. **A.** Schematic representation of the 3TRS method developed in this study. **B.** Representation of the second strand oligo design strategy used to selectively amplify the 3’-end of five individual *HTT/Htt* transcripts. **C.** Unique cDNA read counts of cryptic and canonical human *HTT* and mouse *Htt* transcripts in the three different brain regions analyzed in the WT and the YAC128 HD mouse model. Data are presented as mean + SD. Statistical significance was calculated using an unpaired Student’s t-test: ns (P>0.05), *** (P<0.001), and **** (P<0.0001). **D.** Sashimi plots of WT and YAC128 mouse brain regions showing the reads mapping the expected *HTT* 3’-ends. The arrow indicates the location of the oligonucleotide used for the second strand synthesis. The sequence at the end of the read indicates the polyadenylation signal (PAS) and the cleavage sites in red.

### 3TRS library sequencing and data processing

All 3TRS libraries were sequenced in a NovaSeq X Plus (Illumina) using Novogene pre-made library sequencing service. Sequencing data were analysed using a custom pipeline developed in the laboratory. Quality control of the reads was performed using *FastQC* v0.11.9 (available online at http://www.bioinformatics.babraham.ac.uk/projects/fastqc). After that, samples were trimmed and demultiplexed by barcode with *Cutadapt* v4.2 (31), followed by the extraction of the UMI sequences using *UMI-tools* v1.1.2 (32). Reads were then aligned to the human (GRCh38.p13) or the mouse (GRCm39 M30) genome with *STAR* v2.7.10b, incorporating the gene and transcript annotations from GENCODE release v41 or vM30 with the *STAR* v2.7.10b aligner (33). Only alignments with up to 10 multiple mapping locations were retained, and filtering was applied based on mismatch rate and alignment score relative to read length. Once the BAM files were obtained, duplicated reads were collapsed based on their UMI sequences with *UMI-tools* and indexed with *samtools* v1.16.1 (34). Finally, target regions were quantified using *featureCounts* v2.0.3 (35) from the *Subread* package. Read counts were only considered when the ratio between the raw and UMI collapsed counts showed at least a 10-fold ratio. A *Nextflow* implementation of this workflow is available on *GitHub*, along with all of its containerized dependencies, with extensive information on how to install and run it (https://github.com/blazquezlab/3TRS_analysis_pipeline). The pipeline was implemented within the EPI2ME user-friendly interface and requires four input files: (1) a FASTQ file containing the raw sequencing data, (2) a threeprime.csv file which contains barcode sequences assigned to each sample, (3) a fiveprime.csv file with the information of the individual libraries pooled in each sequencing experiment and (4) a .saf file containing the chromosome coordinates of the target sequences of interest.

### Statistical analysis

Statistical analyses were performed using the GraphPad Prism 10.4.2 software for macOS. Student’s t-test was used when analyzing differences between two groups. Association between variables were tested using the Pearson correlation coefficient (r), which measures linear correlation in data sets. More than two group comparisons were performed with one-way ANOVA followed by multiple comparisons tests. Dunnett’s correction was applied when comparisons were performed against one control group. Tukeýs correction was applied when comparisons were performed between all possible groups. The alpha level for statistical significance was set at < 0.05.

## RESULTS

### *HTT* distal cryptic poly-A site in intron 1 is specifically activated in the YAC128 mouse model upon CAG repeat expansion

To quantify 3’-end processing events, an in-house protocol for 3’-end targeted RNA sequencing (3TRS) was developed. This method was designed to overcome limitations associated with conventional RNA-sequencing approaches and to provide a straightforward, cost-effective workflow for library preparation interrogating pre-defined 3’-ends. The protocol involves three main steps: (1) reverse transcription of each sample independently with a barcoded oligo-dT, (2) a targeted second strand synthesis step in a pool of retrotranscribed (RT) samples with an oligo containing unique molecular identifiers (UMIs) for unique cDNA count, and (3) PCR amplification with oligos containing Illumina adapter sequences followed by sequencing. Raw sequencing data were analyzed using a custom bioinformatic pipeline implemented in Nextflow and executed within the EPI2ME open-source analysis platform. The pipeline generated tab-separated value (TSV) files containing duplicated and deduplicated read counts, quality control reports, and BAM files, which can be easily visualized using genome browsers such as the Integrative Genomics Viewer (IGV) (Fig. 1A). Further methodological details are provided in the Materials and Methods section.

To investigate the effect of CAG repeat expansion in human *HTT* 3’-end processing, 3TRS was performed in brain tissue from YAC128 transgenic mouse model and compared to wild-type (WT) mice (26). This model was selected because it expresses the full-length human mutant *HTT* transgene, thereby enabling the study of human *HTT* 3′-end processing in a relevant *in vivo* context. Importantly, YAC128 mice also retain expression of the endogenous mouse *Htt* gene, allowing the simultaneous investigation of 3′-end processing in both the human and mouse transcripts. To distinguish and quantify the different *HTT* and *Htt* isoforms, second strand specific oligonucleotides were designed to selectively amplify five transcript variants (Fig 1B): (i) two previously reported *HTT* transcripts generated by the aberrant activation of a proximal (hCPA1) or a distal (hCPA2) cryptic pA-sites in the human *HTT* intron 1 (18), (ii) two canonical human *HTT* isoforms, generated by the activation of either a proximal (hPA1) or a distal (hPA2) alternative pA-site in the *HTT* 3’ UTR (36, 37) and (iii) the full-length mouse transcript, which uses the canonical pA-site in the *Htt* 3′ UTR (mPA). Expression of all 3’-ends was measured in the RNA isolated from the striatum (STR; n=4 WT and n=6 YAC), cortex (CTX, n=4 WT and n=7 YAC) and hippocampus (HPC, n=4 WT and n=6 YAC) of YAC128 mice. The endogenous mouse *Htt* transcript, which uses the canonical pA-site (mPA), was detected in all samples, with comparable expression levels between WT and YAC128 mice. Both canonical human *HTT* isoforms, utilizing the proximal (hPA1) and distal (hPA2) pA-sites, were detected exclusively in YAC128 mice carrying the human mutant *HTT* gene. Notably, transcripts using intronic cryptic pA-site were also present only in YAC128 samples. Among these, the distal cryptic pA-site (hCPA2) was consistently detected across all three brain regions analyzed, whereas the proximal site (hCPA1) showed minimal or no expression (Fig. 1C). Sequencing results were further visualized using IGV, which showed the expected sequences from the second strand synthesis oligo to the polyadenylation cleavage site (Fig. 1D, Sup. Fig. 1A). Moreover, 3TRS protocol was specific, as no reads corresponding to other transcripts under study were detected in the libraries prepared with each second strand synthesis oligo (Sup. Fig. 1B). Interestingly, there were no significant differences in the usage of distal cryptic pA-site in human *HTT* (hCPA2) between the different brain regions analyzed, whereas in general a higher expression of canonical human and mouse Huntingtin transcripts was detected in the brain cortex (Sup. Fig. 1C).

### Targeted 3’-end sequencing allows multiplexing several mouse and human Huntingtin transcripts

After quantifying that in human *HTT* gene the distal cryptic poly-A site (hCPA2) is used in the presence of CAG expanded repeats in the YAC128 mouse model, the protocol was optimized to sequence all *HTT/Htt* transcripts simultaneously, so that the number of unique cDNA counts for each pA-site can be compared. To achieve this, second strand cDNA synthesis was performed in a multiplexed reaction combining all second-strand oligonucleotides targeting the predefined pA-sites in the *HTT* and *Htt* genes (Fig. 2A). Importantly, there were no significant differences between the unique cDNA read counts for each target using singleplexed or multiplexed reactions (Sup. Fig. 2A), which demonstrates that multiple oligos can be mixed in the second strand synthesis reaction to simplify the protocol and get comparable reads of each interrogated pA-site.

**Figure 2:**
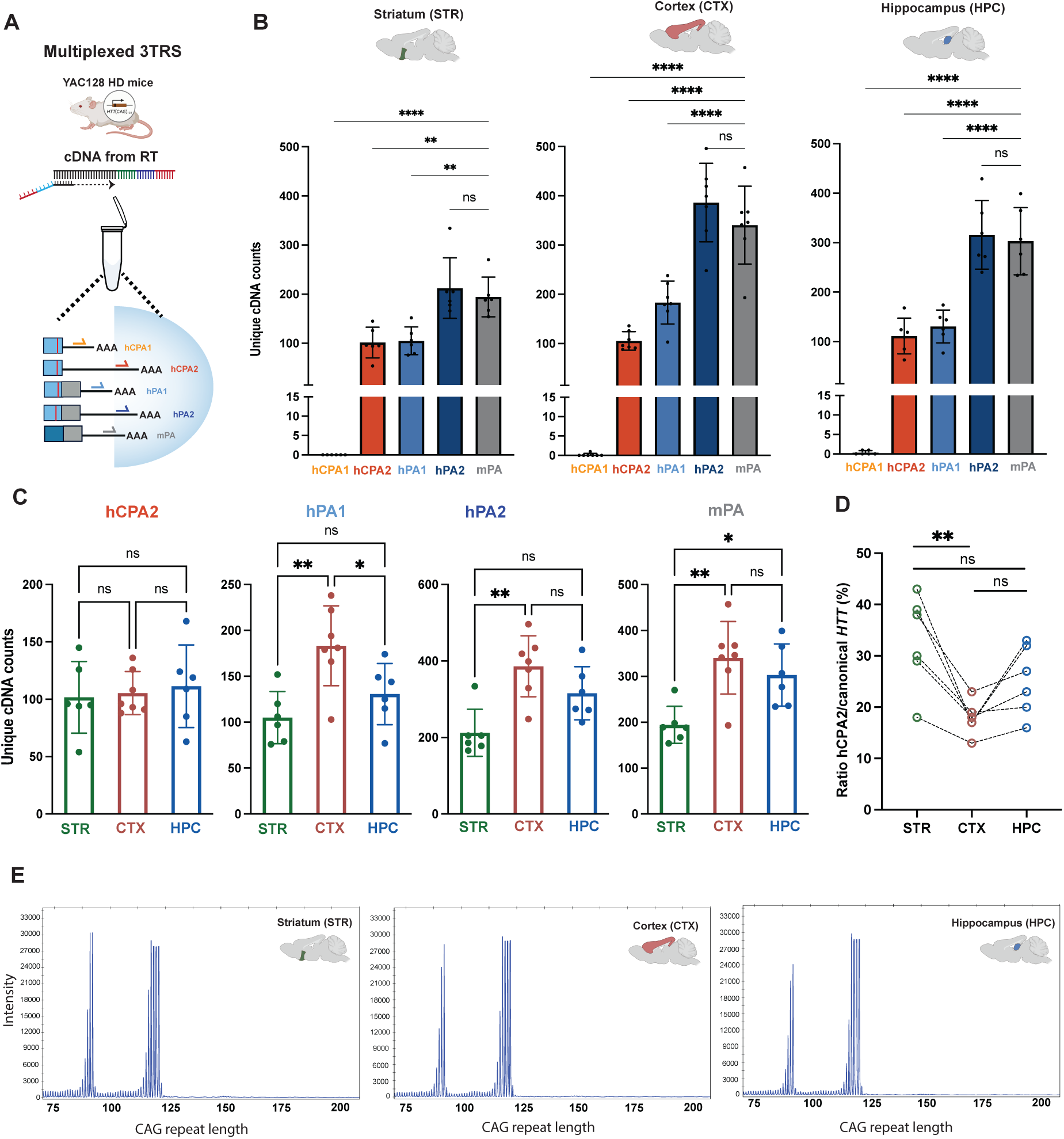
Multiplexed 3TRS in the YAC128 mouse model. **A.** Representation of the second strand synthesis reaction multiplexing all oligonucleotides targeting the predefined poly-A sites in the *HTT*/*Htt* transcripts. **B.** Unique cDNA read counts for all interrogated *HTT/Htt* 3’-ends across three brain regions. Data are presented as mean + SD. Statistical significance was calculated with a one-way ANOVA test followed by Dunnett’s multiple comparisons analysis. **C.** Unique cDNA read counts of human *HTT* and mouse *Htt* transcripts in the three different brain regions analyzed. Data are presented as mean + SD. Statistical significance was calculated with a one-way ANOVA test followed by Tukeýs multiple comparisons analysis. **D.** Ratio of cryptic hCPA2 versus all canonical *HTT* isoforms in the striatum (STR), cortex (CTX) and hippocampus (HPC) of YAC128 transgenic mice. Data are presented as individual values. Statistical significance was calculated using a one-way ANOVA test followed by Tukeýs multiple comparisons analysis. **E.** Length distribution of *HTT* CAG tract in the striatum, cortex or hippocampus of one representative YAC128 mouse. Across the figure, statistical significance is indicated as ns (P > 0.05), * (P ≤ 0.05), ** (P ≤ 0.01), *** (P ≤ 0.001) and **** (P ≤ 0.0001).

Across all three brain regions, activation of distal hCPA2 cryptic pA-site, but not proximal hCPA1, was confirmed in YAC128 mice (Fig. 2B). In multiplexed 3TRS aligned sequencing reads were visualized from a single BAM file which contains the 3’-end fragments from all the interrogated human pA-sites (Sup. Fig. 2B). As in the individual reactions, no significant differences in hCPA2 unique cDNA counts were observed between the different brain regions analyzed (Fig. 2C). Since a higher level of canonical human *HTT* and mouse *Htt* was present in the cortex, cryptic *HTT* expression level was calculated relative to all canonical *HTT* isoforms. This analysis showed that the ratio of cryptic hCPA2 versus canonical *HTT* was significantly higher in the striatum compared to cortex (Fig. 2D). *HTT* fragment analysis confirmed that the YAC128 brain shows the expected repeat length distribution, with no appreciable somatic repeat instability (Fig. 2E).

### Both *Htt* cryptic poly-A sites are used in *Hdh^Q111/+^* knock-in mice

In contrast to YAC128 mice, the *Hdh^Q111/+^* knock-in (KI) mouse model carries a chimeric human/mouse exon 1 with a 111 poly-Q tract within the endogenous mouse *Htt* gene (38) (Fig. 3A). Therefore, the CAG expansion occurs in the *Htt* genomic context, resembling more closely HD patients. Previous studies in HD KI mice have described two cryptic poly-A sites in *Htt* intron 1 (mCPA1 and mCPA2), which differ from the human *HTT* cryptic poly-A sites. These sites are located at positions 680 bp and 1145 bp in *Htt* intron 1, respectively (Fig. 3A) (18). In order to investigate activation of cryptic polyadenylation in the mouse *Htt* gene upon CAG repeat expansion, RNA samples from the brain cortex (CTX), hippocampus (HPC), and striatum (STR) of WT (N=4) and *Hdh^Q111/+^* (N=3) mice were analyzed using 3TRS. As in the YAC128 model, second strand synthesis oligos were designed to selectively amplify three transcript variants (Fig 3B): (i) the mouse canonical poly-A site (mPA), arising from the wild-type and the mutant forms, (ii) the mouse proximal cryptic poly-A site (mCPA1) and (iii) the mouse distal cryptic poly-A site (mCPA2). Based on previous results in the YAC128 model, all second strand oligos were multiplexed in a single reaction (Fig. 3B).

**Figure 3:**
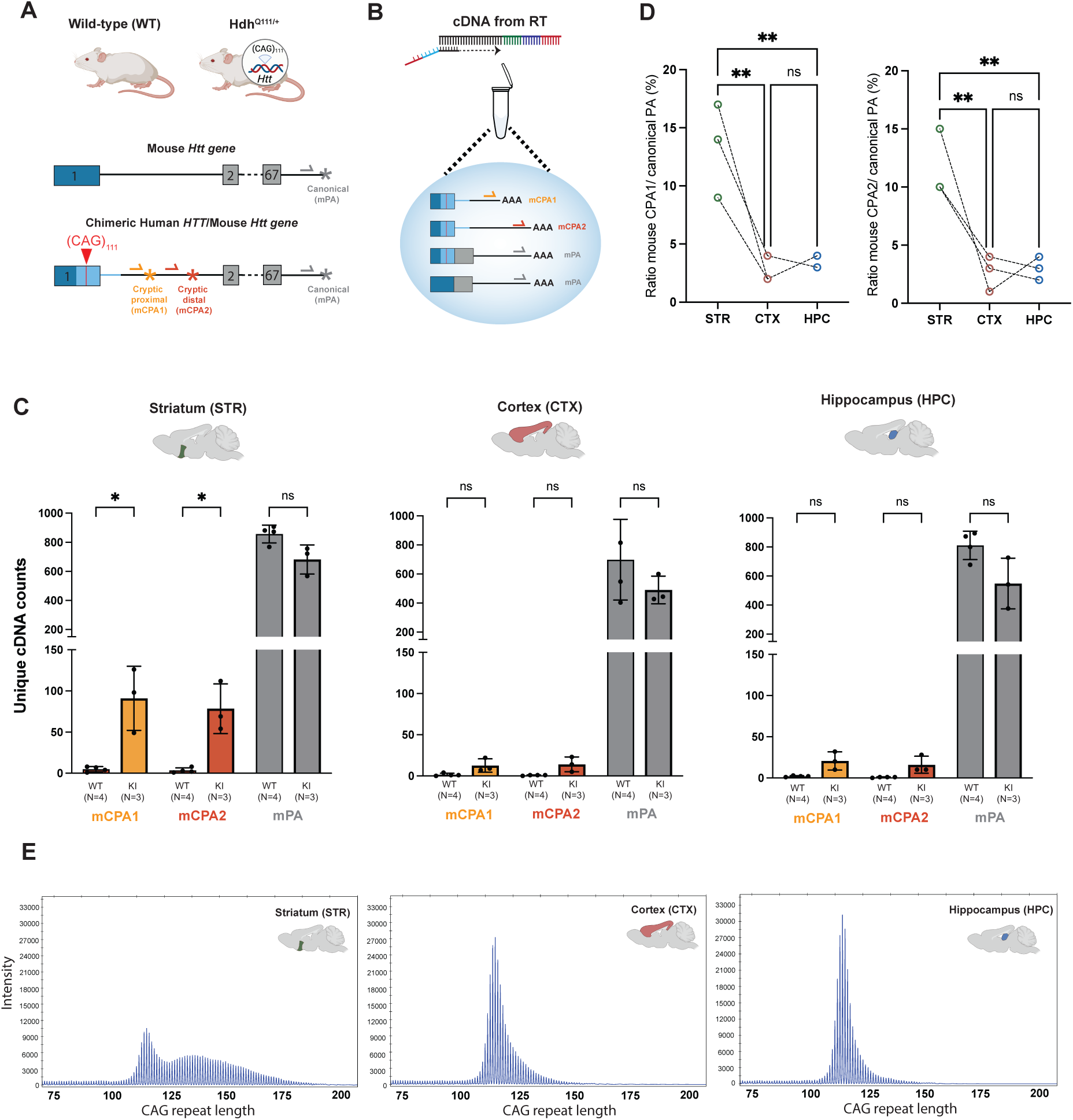
*Htt* RNA processing in the *HdhQ111/+* mouse model. **A.** Representation of the second strand oligo design strategy used to selectively amplify the 3’-end of three mouse *Htt* transcripts. **B.** Representation of the second strand synthesis reaction multiplexing all oligonucleotides targeting the predefined poly-A sites in the *Htt* transcripts. **C.** Unique cDNA read counts of cryptic and canonical mouse *Htt* transcripts in the three different brain regions analyzed in the WT and the *Hdh^Q111**/+**^* HD mouse model. Data are presented as mean + SD. Statistical significance was calculated using an unpaired Student’s t-test. **D.** Ratio of cryptic versus canonical *Htt* isoforms in the striatum (STR), cortex (CTX) and hippocampus (HPC) of *Hdh^Q111**/+**^* mice. Data are presented as individual values. Statistical significance was calculated with a one-way ANOVA test followed by Tukeýs multiple comparisons analysis. **E.** Length distribution of *Htt* CAG tract in the striatum, cortex or hippocampus of one representative Hdh^Q111**/+**^ mouse. Across the figure, statistical significance is indicated as ns (P > 0.05), * (P ≤ 0.05), and ** (P ≤ 0.01).

Compared to the YAC128 mouse model, sequencing results showed that both *Htt* mCPA1 and mCPA2 sites were activated in *Hdh^Q111/+^* mice upon CAG repeat expansion. Moreover, the KI mice exhibited the highest number of cryptically polyadenylated transcripts in the striatum, whereas the differences obtained in the cortex and the hippocampus of *Hdh^Q111/+^* mice compared to wild-type did not reach statistical significance (Fig. 3C). When the expression level of each cryptic transcript was corrected against canonical *Htt* expression, we confirmed that cryptic polyadenylation was significantly higher in the striatum (Fig. 3D). Since previous studies have reported that the CAG repeat expansion undergoes tissue-specific somatic instability, most prominently in the striatum (39), we investigated whether the highest activation of cryptic polyadenylation observed in the striatum was associated with the level of somatic instability in that tissue. Indeed, *Htt* fragment analysis showed that cryptic polyadenylation increases upon somatic repeat expansion in the striatum (Fig. 3E). Aligned sequencing reads were visualized in IGV using the mouse genome as reference, which confirmed that cryptic polyadenylation in *Htt* intron 1 is activated in the striatum of *Hdh^Q111/+^* mice, whereas canonical transcripts were present in all tissues from wild-type and *Hdh^Q111/+^* mice (Sup. Fig. 3B).

### *HTT* cryptic polyadenylation occurs in genetically edited HEK293 HD cell lines

To translate the findings from HD mouse models into a human cellular context, we studied HEK293 cell lines carrying homozygous CAG repeat expansions, with or without CAA interruptions, in the endogenous *HTT* locus. These include seven previously published cell lines with homozygous CAG expansions (41/41, N= 1, 53/53; N=2, 83/83; N= 4) (21), and three unpublished lines homozygous for a 41 polyQ tract but including CAA interruptions (39CAG-2XCAA; N= 4, 40CAG-1XCAA; N=1) along with four wild-type cell lines (WT; N=4) (Fig. 4A). Following the same approach as before, a multiplexed library targeting all human *HTT* transcripts (hCPA1, hCPA2, hPA1 and hPA2), was prepared and sequenced (Fig. 4B).

**Figure 4:**
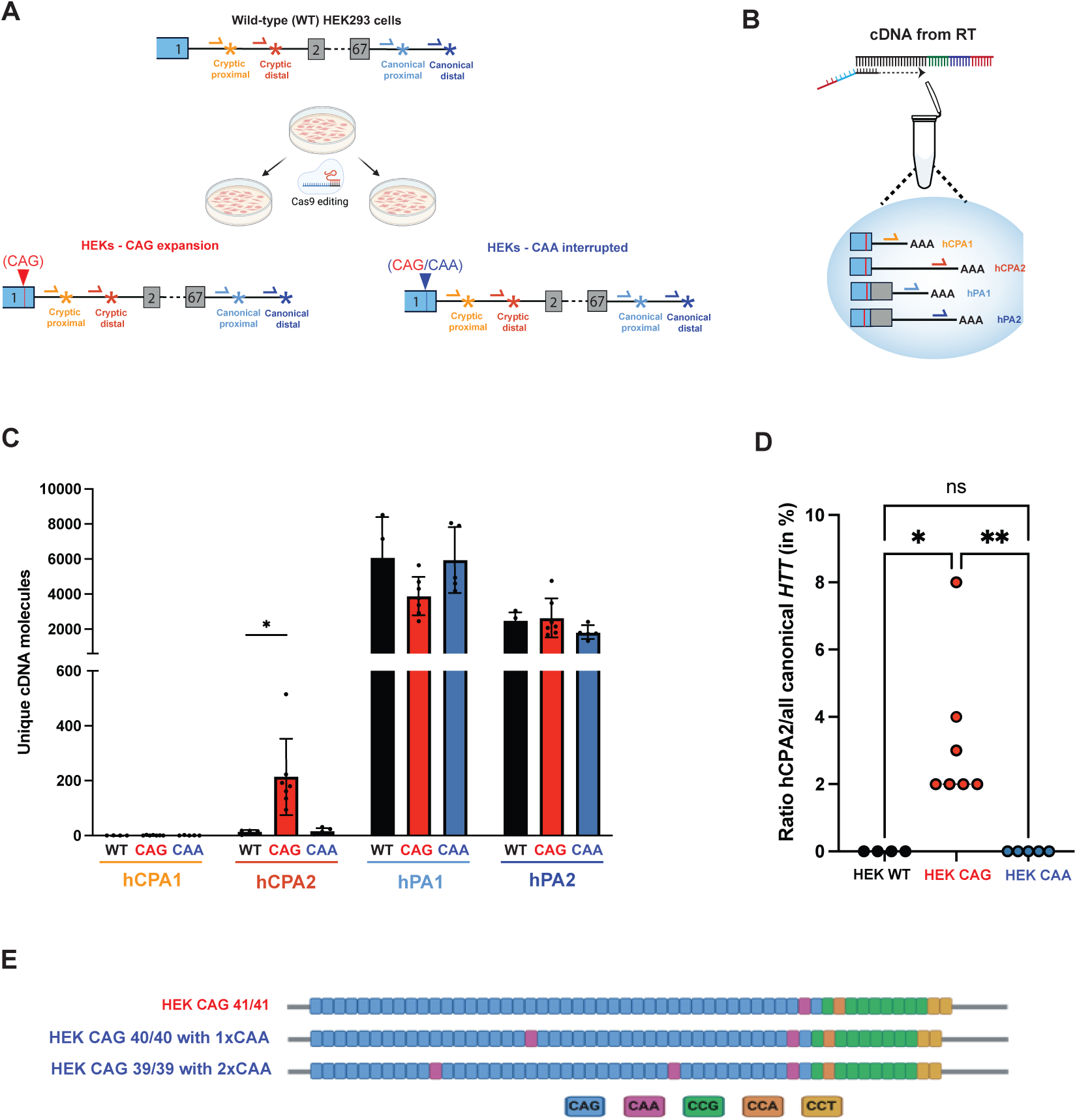
*HTT* cryptic polyadenylation in HEK293 HD cell lines. **A.** Image of the *HTT* gene in the three HEK293 cell lines analyzed in this study, highlighting the second-strand oligos used to amplify the 3’-ends of proximal and distal cryptic and canonical *HTT* transcripts. **B.** Representation of the second strand synthesis reaction multiplexing second-strand oligonucleotides targeting all *HTT* transcripts. **C.** Unique cDNA read counts of cryptic and canonical *HTT* transcripts in the three HEK293 cell lines under study. Data are presented as mean + SD. For each *HTT* transcript, statistical significance was calculated using a one-way ANOVA test followed by Dunnett’s multiple comparisons analysis. Only statistically significant results are indicated as * (P ≤ 0.05). **D.** Ratio of cryptic versus canonical *HTT* isoforms in the three investigated HEK293 cell lines. Data are presented as individual values. Statistical significance is indicated as ns (P > 0.05), * (P < 0.05) and ** (P ≤ 0.01), after calculation with a one-way ANOVA test followed by Tukeýs multiple comparisons analysis. **E.** Motif and length configuration of *HTT* CAG tract in one representative HEK-CAG cell line and two HEK-CAA cell lines.

Regarding canonical *HTT* expression, no significant changes in the proximal (hPA1) or the distal (hPA2) poly-A sites were observed between WT, CAG-expanded and CAA-interrupted HEK293 cell lines (Fig. 4C). Interestingly, while in the YAC128 HD mouse model the distal canonical poly-A site (hPA2) was more frequently used than the proximal one (hPA1), in HEK293 HD cells the proximal site was more active. These findings align with previous reports indicating that the longer isoform (hPA2) is more abundant in brain tissue, whereas the shorter isoform (hPA1) predominates in peripheral tissues (37). Concerning cryptic polyadenylation, the distal site (hCPA2), but not the proximal one (hCPA1) was again only used in HEK293 cells carrying homozygous CAG repeat expansions (N=7). Remarkably, CAA interruptions (N=4) avoided *HTT* cryptic poly-A site usage, to a similar extent as wild-type HEK293 cells (N=4), which also did not show cryptic polyadenylation (Fig. 4C). When the expression level of cryptic *HTT* was corrected with canonical *HTT* expression, it confirmed that cryptic *HTT* expression occurs exclusively in the context of pure CAG repeat tracts (Fig. 4D). These findings suggest that the presence of CAA interruptions may prevent the aberrant activation of cryptic poly-A sites in the context of expanded CAG repeats, potentially contributing to their protective effect in HD phenotype (40).

In CAG-expanded HEK293 cells repeat length did not correlate with the *HTT1a* level (Sup. Fig. 4A). Long-read sequencing of *HTT* CAG-repeat fragment confirmed the expected motif distribution (Fig 4E). Aligned sequencing reads were visualized in IGV, which confirmed that *HTT* cryptic polyadenylation was exclusive from HEK293-CAG HD cells, whereas both canonical sites were recognized across all the three different cell lines (WT, CAA, and CAG) (Sup. Fig. 4B).

### *HTT* cryptic polyadenylation in HD patient-derived samples requires ultralong CAG repeat expansions

To assess *HTT* cryptic polyadenylation in HD patient-derived samples, we performed 3TRS in primary fibroblasts derived from control (WT, N = 8) and HD patients (HD, N = 16). Control samples contained less than 40 CAG repeats in the *HTT* gene. HD patient-derived fibroblasts were divided in four groups according to their CAG-repeat size: CAG 40-50 (N=13), CAG 56 (N=1), CAG 81 (N=1) and CAG 180 (N=1). Library preparation was performed multiplexing second-strand oligos targeting all cryptic and canonical proximal and distal poly-A sites (Fig. 5A). In control and HD fibroblasts, reads mapping the canonical proximal poly-A site were significantly more abundant than reads mapping the distal site, in accordance with the results obtained in the non-neural HEK293 cell line. Canonical *HTT* expression showed no differences between control and CAG 40-50 groups. Regarding cryptic site activation, there were no reads mapping the proximal cryptic poly-A site (hCPA1) in none of the samples, and reads mapping the distal site (hCPA2) were only present in the fibroblast cell line with 180 CAG repeats (Fig. 5B), meaning that ultra-long CAG repeats are required to activate the distal cryptic poly-A site in human fibroblasts. With 3TRS, the unique cDNA reads mapping each of the interrogated 3’-ends can be compared against each other. This analysis revealed that cryptic *HTT* transcripts represent around 5% of all canonical *HTT* transcripts, only in the CAG-180 fibroblast cell line (Fig. 5C). *HTT* fragment analysis confirmed the expected repeat length distribution, shown for three lines with the longest CAG repeat expansions (Sup. Fig. 5A).

**Figure 5:**
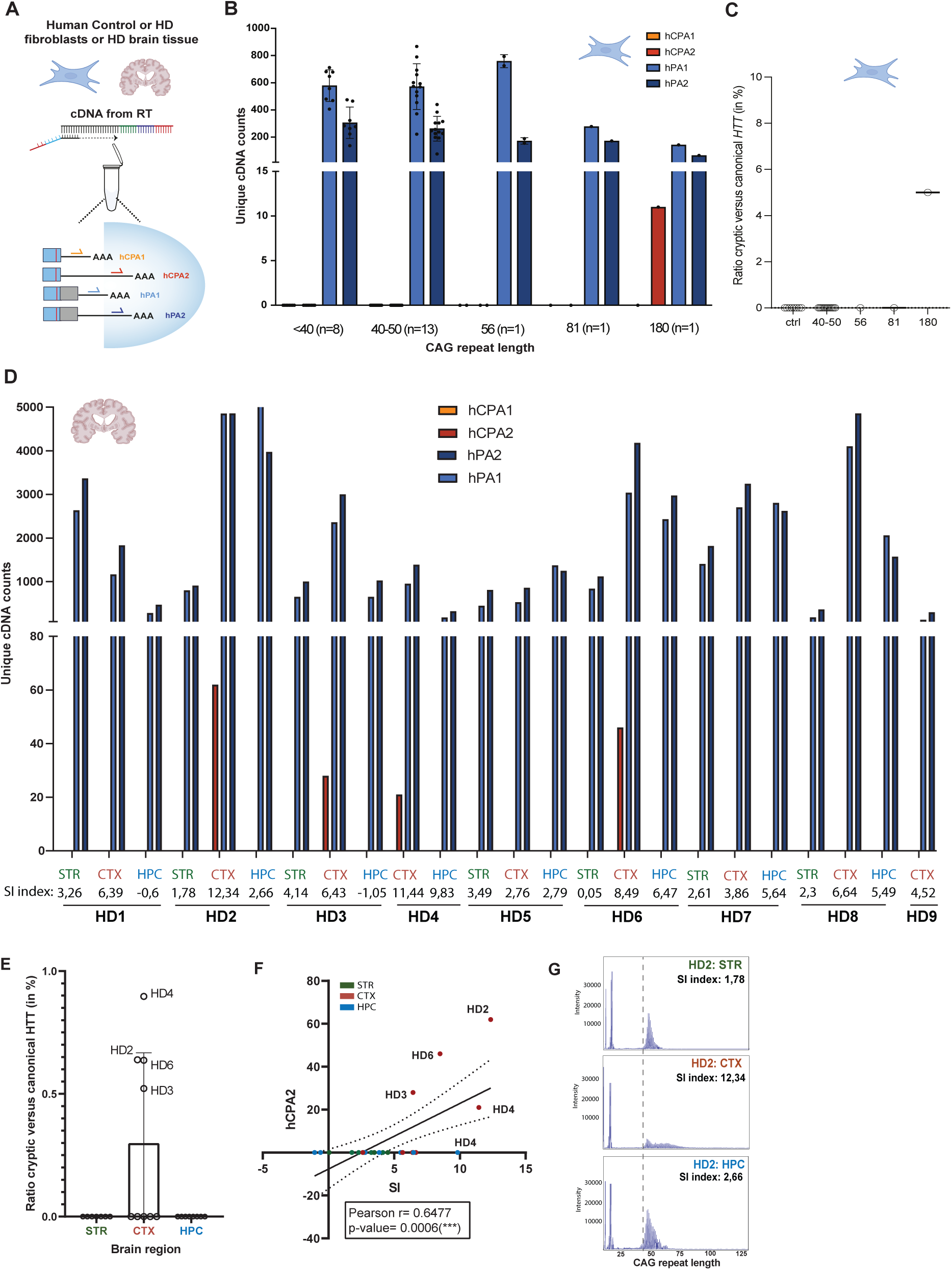
*HTT* cryptic polyadenylation in human HD samples. **A.** Representation of the second strand synthesis reaction multiplexing four second-strand oligonucleotides targeting *HTT* transcripts, starting from RNA isolated from fibroblasts or postmortem brain tissue. **B.** Unique cDNA read counts of cryptic and canonical *HTT* transcripts in human fibroblasts. Data are presented as mean + SD, whenever the groups include more than one sample. Statistical significance was calculated using an unpaired Student’s t-test: ns (P>0.05) and * (P<0.05). **C.** Ratio of cryptic versus canonical *HTT* isoforms in control and HD fibroblasts. **D.** Unique cDNA read counts of cryptic and canonical *HTT* transcripts in the human HD brain regions under study starting with 4 μg RNA. Data are represented as individual values. **E.** Ratio of cryptic versus canonical *HTT* isoforms in HD post-mortem brain samples. **F.** Correlation analysis between somatic instability index and hCPA2 counts from the postmortem samples. Association between somatic instability index and hCPA2 expression was tested using the Pearson correlation coefficient (r). **G.** Length distribution of *HTT* CAG tract in the postmortem striatum, cortex and hippocampus of one representative HD patient. The vertical dashed line sets the 40 CAG repeat length threshold. The somatic instability index (SI) is indicated for each sample.

Finally, postmortem brain tissue from HD patients was analyzed. Importantly, in parallel to 3TRS library preparation, we calculated the somatic instability (SI) index starting from the same RNA aliquot, as explained in the Material and Methods section. In a first attempt, 1μg RNA was used for library preparation, isolated from the bulk cortex (CTX, N=5), hippocampus (HPC, N=2), and striatum (STR, N=2) of five different HD patients. Using the 3TRS strategy described above for human fibroblasts (Fig. 5A), canonical *HTT* transcripts were detected in all brain samples (Sup Fig. 5B), where a higher abundance of the canonical distal poly-A site was observed compared to human fibroblasts. Regarding cryptic polyadenylation, our results show that only the distal poly-A is active, but not in all HD brain patients and regions. Instead, cryptic polyadenylation was only detected in those brain regions with the highest SI indexes (Sup Fig. 5B).

To increase the detection of the very low levels of *HTT1a* transcripts, we repeated 3TRS increasing the starting amount of total RNA to prepare the 3TRS library from 1ug to 4ug. This required the extraction of a new RNA aliquot from all HD1 brain regions. Moreover, 3TRS with 4ug of RNA as starting material was performed in additional brain regions from HD2, HD3 and HD4 cases and in additional HD cases (HD5-HD9). Unfortunately, no more RNA or tissue was left from HD10 to repeat 3TRS library with a 4ug input. Altogether, 24 HD postmortem samples were analysed, isolated from cortex (CTX, N=9), hippocampus (HPC, N=8), and striatum (STR, N=7) from 9 HD cases. Canonical *HTT* transcripts were detected in all brain samples (Fig. 5D), while the distal cryptic polyadenylation event leading to the *HTT1a* transcript was only detected in four HD brain samples (Fig. 5D). Contrary to our initial expectations, *HTT1a* reads were captured only in the cortex and not in the striatum, despite the latter being the most vulnerable brain region in HD. In agreement with this result, the SI indexes obtained in HD striatum were very low, likely reflecting the extensive neuronal loss present in this tissue in late-stage HD patients. Overall, cryptic *HTT* transcripts represent less than 1% of all canonical *HTT* transcripts (Fig. 5E). Remarkably, a statistically significant correlation was observed between the number of cryptic *HTT* reads and the SI index (Fig. 5F).

We also noted that not only the SI index but also the inherited CAG repeat-length (Sup. Table 1) may influence cryptic *HTT1a* expression. This was observed in HD3 and HD8 cortex samples, where reads mapping the cryptic distal site (hCPA2) were only captured in HD3 cortex despite having a lower SI index than HD8 cortex (6, 43 versus 6, 64) (Fig. 5D, Sup. Fig. 5C). In HD3 patient, however, the initial CAG repeat length in the mutant *HTT* allele, as genotyped in peripheral blood, was longer (54 CAG repeats) than in HD8 (45 CAG repeats). Another interesting observation was that in HD1 results regarding *HTT1a* expression differ between two independent RNA aliquots isolated from the same brain region. In the first RNA isolation, cryptic *HTT* reads were present in HD1 striatum, which exhibited a SI index of 9, 87 (Sup. Fig. 5B). In a second RNA isolation, performed later starting from the same initial tissue block to increase the RNA amount available to repeat 3TRS with a 4ug RNA input, no cryptic reads were detected (Fig. 5D). This negative result was consistent with the low SI index reported in the second RNA aliquot (Sup. Fig. 5C). Finally, in the hippocampus, none of the samples showed *HTT1a* expression, not even in the presence of a high SI index, as was the case for HD4 patient (Fig. 5D). In summary, *HTT1a* expression was primarily detected in cortical samples with high SI indexes, as shown for HD2 patient as an illustrative example (Fig. 5G).

## DISCUSSION

In this work we have established a targeted RNA-sequencing strategy to investigate the simultaneous expression of several *HTT* transcripts generated by canonical and cryptic polyadenylation in the context of HD. Our results agree with previous studies that initially described *HTT1a* expression by incomplete *HTT* splicing in the presence of expanded CAG repeats (18–20, 41). Compared to previous studies however, which primarily detect *HTT* intron 1 fragments by RT-PCR (21, 22), quantitative PCR (qPCR) (18, 19), digital PCR (20) or probe based-assays (17) as a proxy for *HTT1a* expression, the strategy presented here involves RNA sequencing of the 3’-ends of polyadenylated *HTT* transcripts. Our approach (i) provides quantitative information from 3’-end processed mRNA isoforms and avoids quantification of yet unprocessed *HTT* intron 1 molecules; (ii) precisely differentiates between alternative isoforms generated by the usage of nearby poly-A sites; (iii) allows multiplexing of dozens of samples early in the library preparation protocol, adding a sample barcoding step, and (iv) avoids PCR-based amplification biases introducing Unique Molecular Identifiers (UMIs) during library preparation. Finally, the protocol allows multiplexing several oligos in the second strand synthesis reaction, which allows comparing absolute values from alternative isoforms, also a limitation from previously used strategies (17, 19, 20).

Cryptic polyadenylation in *HTT*/*Htt* was first described by 3’RACE (18), a method that confirmed the usage of a poly-A site, but not in a quantitative-manner. Follow-up studies developed quantitative assays using amplification primers or fluorescent probes, such as qPCR (19), digital PCR (20) and a QuantiGene probe-based assay to measure *HTT1a* expression (17). These assays, however, do not rely on RNA-sequencing and cannot precisely map and quantify reads to specific polyadenylation sites with *Huntingtin* transcripts. Indeed, primers or probes that bind proximal *HTT* intron 1 sequences cannot differentiate between transcripts that use the proximal or the distal poly-A sites, because all distal transcripts contain the proximal intron 1 sequences. Our results using 3TRS clearly demonstrate that in YAC128 transgenic mice, as in all human samples evaluated in this study, *HTT1a* transcript derives exclusively from the activation of the distal cryptic poly-A site (hCPA2). Although a slight, but not significant, increase in the number of reads derived from the proximal cryptic site (hCPA1) was detected when performing singleplexed 3TRS in the YAC128 mouse model, this difference was no longer present when all second strand oligos were multiplexed, which allows the comparison of absolute values from alternative isoforms. *HTT1a* expression also shows species-specific differences, since in the *Hdh ^Q111/+^* knock-in mice model, which contains the mouse *Htt* intron 1 sequence downstream the expanded CAG repeat, both proximal and distal poly-A sites are used.

Our results also have important implications regarding HD pathogenicity. Using 3TRS we demonstrate that activation of *HTT* cryptic polyadenylation occurs in mouse and human brain, in a mechanism that is exacerbated upon the somatic CAG-repeat instability that occurs in specific regions of the brain. In YAC128 mice, *HTT* hCPA2 expression was comparable across the striatum, cortex and hippocampus. This could be due to the mutant allele being in a genetically engineered chromosome, or to the presence of interspersed CAA codons along the exon 1 polyQ tract that stabilize the CAG repeat and prevent germline and somatic instability (42, 43). Compared to cortex, however, the striatum showed a slight but significantly higher ratio of cryptic versus canonical *HTT*, which may contribute to the selective striatal neuronal loss reported in this model (26). In *Hdh^Q111/+^* mice, the striatum clearly showed the highest level of proximal and distal cryptic *HTT*, a finding that correlated with a higher somatic instability in this brain region and may explain the stronger vulnerability of the striatum in this HD model (44, 45). These results underscore the relevance of using HD models where CAG repeat expansions are introduced within the endogenous locus to investigate HD pathology.

The data obtained in HEK293-CAG and CAA cell lines support a model where pure CAG repeats are required to activate *HTT* cryptic polyadenylation and *HTT1a* expression. Previous studies have shown that the presence of CAA interruptions within the CAG repeat tract of the *HTT* gene are associated with a milder HD phenotype (40, 46). Our results suggest that CAA interruptions may exert a protective effect by modifying the regulatory landscape of *HTT* cryptic polyadenylation, reducing *HTT1a* expression. The results obtained in these cell lines, however, should be interpreted with caution. HEK293-CAG lines are homozygous for CAG repeat expansion which could explain why cryptic polyadenylation is detected with only 41 CAG repeats, and without a positive correlation with increasing CAG repeat lengths. We cannot either disregard that the genetic background, the limited number of samples or the non-neural origin of HEK293 cell line influences these results. In HD patient-derived samples, our results indicate that ultra long CAG repeat expansions are required for the activation of *HTT* cryptic poly-A sites. In HD fibroblasts, *HTT1a* expression was detected only in a cell line derived from a juvenile-onset HD patient with 180 CAG repeats. These findings are consistent with previous studies (19, 20), but the use of 3TRS offers additional advantages. First, it enables a direct comparison of cryptic versus canonical read counts, revealing that approximately 5% of all *HTT* transcripts in this cell line correspond to the toxic *HTT1a* isoform. Second, the incorporation of UMIs avoids and corrects PCR amplification bias, and reveals that *HTT1a* transcripts arise from a very small number of unique RNA molecules that generate a disproportionately high number of duplicated sequencing reads.

Finally, analysis of human postmortem brain tissue demonstrates that *HTT* distal cryptic poly-A site is also activated in HD brain upon CAG repeat expansion. Notably, cryptic *HTT* expression was barely detected in the striatal samples we have analysed, but instead in the cortical samples exhibiting the highest somatic instability (SI) indexes. These results likely reflect the extreme atrophy that characterizes the striatum in HD, where ∼95% of the medium spiny neurons (MSNs) are lost in advanced disease (47). Indeed, RNA samples from bulk HD striatum showed low SI indexes, consistent with the idea that the somatically expanded MSNs expected in this region have already been lost (3, 5). *HTT1a* expression was detected in the striatum of a single HD patient (HD1), the only sample with a high SI index. In contrast, *HTT1a* reads were not detected in a second RNA aliquot isolated from the same tissue block, which showed a much lower SI index. This discordance likely reflects substantial spatial heterogeneity in MSN loss within the same region of a given HD patient, a well-known feature of HD pathology (48). In human HD brain, *HTT1a* expression was almost exclusively reported in the cortex, where HD pathology appears later in time (47) and CAG somatic mosaicism has been recently reported in specific cortical neurons (4, 49, 50).

Importantly, a high SI index does not necessarily translate into detectable *HTT1a* expression. The initial germline CAG repeat length may modulate the SI threshold at which cryptic *HTT1a* transcripts emerge in the brain, such that individuals starting with longer CAG tracts, such as HD3, may reach this threshold at lower levels of somatic expansion. In addition, tissues may differ in how SI contributes to *HTT1a* expression. For instance, in the hippocampus we did not detect cryptic reads supporting *HTT1a* expression, despite the high SI index reported in one HD patient (HD4, SI index: 9, 83), suggesting that somatic expansion alone is insufficient to drive *HTT1a* expression across all brain regions.

Future studies that apply 3TRS to additional HD samples and starting from RNA isolated from single cells or cell subpopulations rather than bulk tissue will help us understand the complex interplay between somatic repeat expansion and *HTT* cryptic polyadenylation. In the meantime, our results support a model where long-somatic DNA repeat expansions acquired through aging drive neurodegeneration in HD, promoting *HTT* cryptic polyadenylation, *HTT1a* RNA and protein expression and disease pathology, in line with recently published studies (2, 5, 51).

This study also has important implications for the development of therapeutic strategies targeting *HTT1a* isoform with antisense RNA molecules (52, 53). The successful evaluation of ongoing and future therapies requires quantitative, sensitive, and cost-effective RNA-sequencing-based methods which can simultaneously evaluate the expression of all *HTT* isoforms using standardized protocols that allow multiplexing several samples. In addition, 3TRS could help elucidate the mechanisms driving *HTT1a* formation, which are yet not fully understood. Several studies point to a complex scenario involving splicing factors, RNA polymerase II transcriptional speed, and epitranscriptomic modifications (28, 54, 55). Our approach also captures physiological switches in alternative poly-A site usage. According to our data, in human cells the short *HTT* 3’UTR isoform (hPA1; 10.3kb) predominates in proliferative conditions, as observed in HEK293 cells and fibroblasts, whereas the long isoform (hPA2; 13.7kb) is more abundant in brain, as previously reported (36).

The implications of aberrant 3′-end processing in human disease remain poorly understood, partly due to technical challenges to detect alternative and cryptic 3’-ends from standard short-read RNA-sequencing data (56). The 3TRS approach presented here can be readily adapted to quantify multiple transcripts generated by alternative polyadenylation across diverse physiological and/or disease scenarios. We provide comprehensive experimental details and a standardized analysis pipeline that requires minimal coding and computational expertise, enabling a straightforward implementation of this method.

In summary, in this work we establish a targeted 3’-end RNA sequencing method (3TRS) to quantitatively measure *HTT* transcripts generated by cryptic polyadenylation in HD. Our results demonstrate that activation of a distal cryptic polyadenylation site in *HTT* intron 1 produces the pathogenic *HTT1a* transcript, particularly in the presence of long, uninterrupted CAG repeat expansions. In HTT/Htt chimeric mouse models, cryptic polyadenylation is enhanced in brain regions with higher somatic CAG instability and neuronal vulnerability, such as the striatum. In human cells and patient samples, *HTT1a* expression requires very long CAG tracts and is blocked by CAA interruptions, supporting a model in which pure, expanded repeats drive toxicity. In postmortem HD brain, *HTT1a* levels reflect somatic instability but not the striatal vulnerability, likely due to the end-stage of the HD patients included in this study. These findings support a disease mechanism in which age-dependent somatic repeat expansion promotes *HTT1a* production and provide a sensitive sequencing-based tool for evaluating *HTT*-targeting therapies.

## ACKNOWLEDGEMENTS

We are grateful to HD patients and their families for donating biological samples for research. We would like to acknowledge Prof. Gill Bates and Dr. Sandra Fienko (Institute of Neurology, UCL, London) for their initial involvement providing RNA samples that were not included in this work, and Dr. Sonia Vazquez-Sanchez, Dr. Carlos Marinas-Chillón and Dr. Jone López-Erauskin, from Prof. Don Cleveland’s group (UCSD, USA) for critical discussion of this work. We want to acknowledge all members of the ‘Neurogenetics, RNA biology and therapies’ group at Biogipuzkoa Health Research Institute for shared discussion. We finally want to thank the DIPC Atlas Supercomputing Center and their support team for data storage and technical assistance with computational data analysis.

## AUTHOR CONTRIBUTIONS

LB designed and supervised the study in collaboration with AVB, VB, JU, MH, MO and ALM. AVB, MM and OAG performed experiments. AHR, MCH, AVB, MM and LB analyzed data. AM generated the HEK293-CAA modified cell lines and EPN collected animal tissue. AR collected and clinically characterized human postmortem brain tissue. LB oversaw project administration and provided supervision. The initial draft was written by LB and AVB and all authors contributed to review and editing of the manuscript.

## SUPPLEMENTARY DATA

Supplementary Data are available at NAR online.

## DATA AVAILABILITY

The original and processed sequencing data generated in this study are available from the corresponding author upon request. The bioinformatic pipeline developed for 3TRS analysis is already available via GitHub, and methodological details are described comprehensively to ensure reproducibility.

