## Supplementary Figures for "Targeted 3’-end RNA sequencing uncovers cryptic polyadenylation in Huntington’s disease linked to somatic instability and CAG repeat purity"

#### Supplementary Figure 1

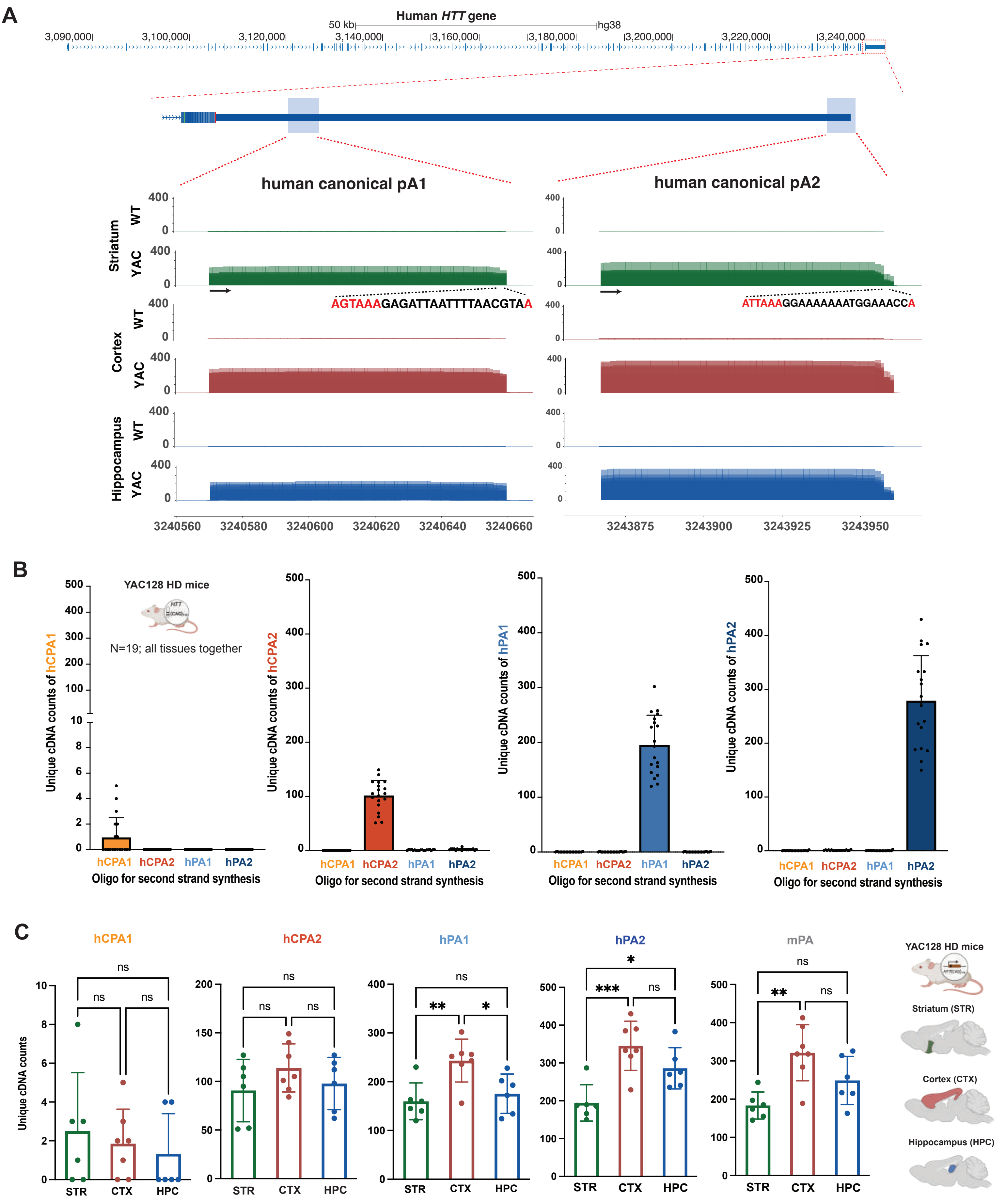

### Supplementary Figure 2

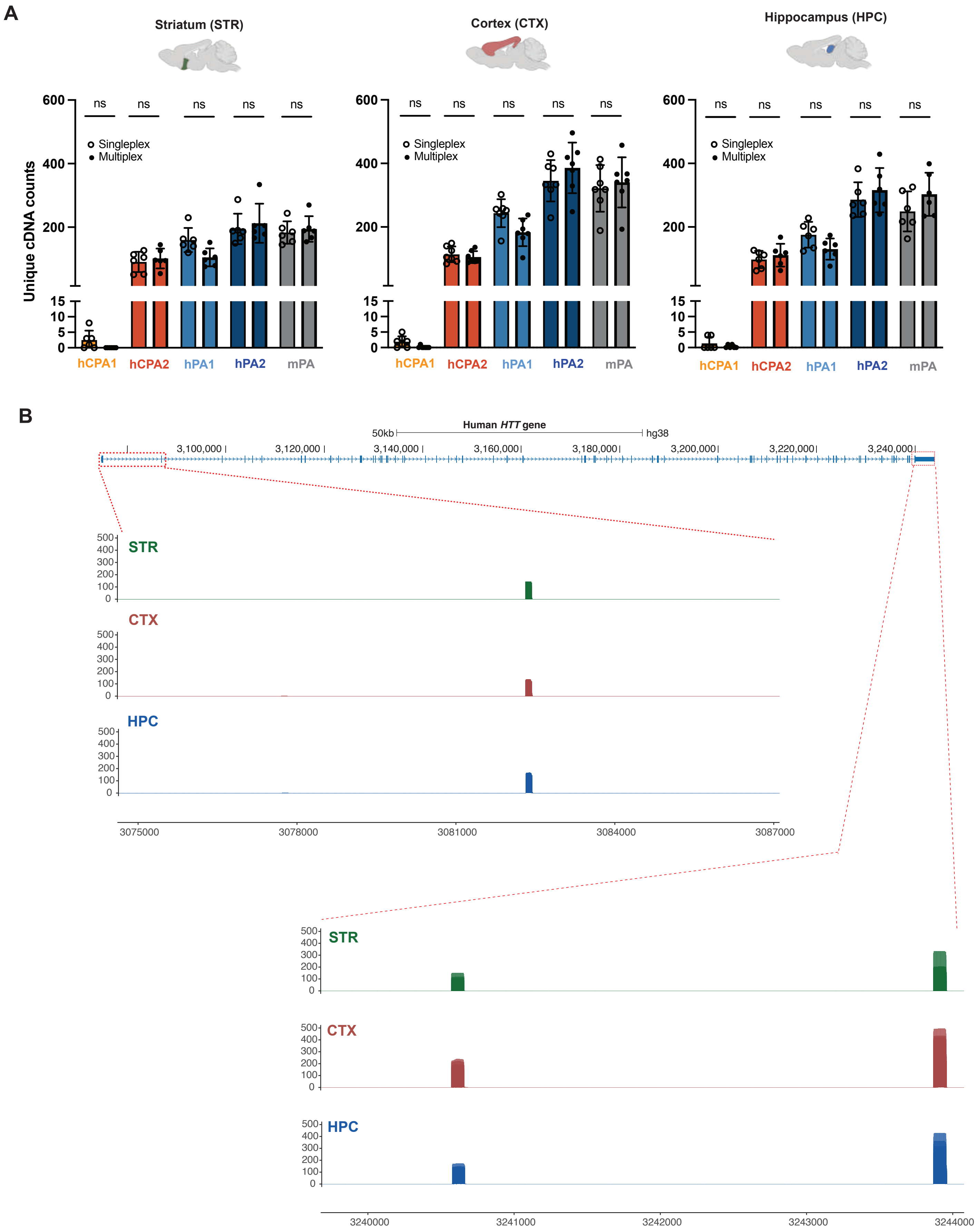

### Supplementary Figure 3

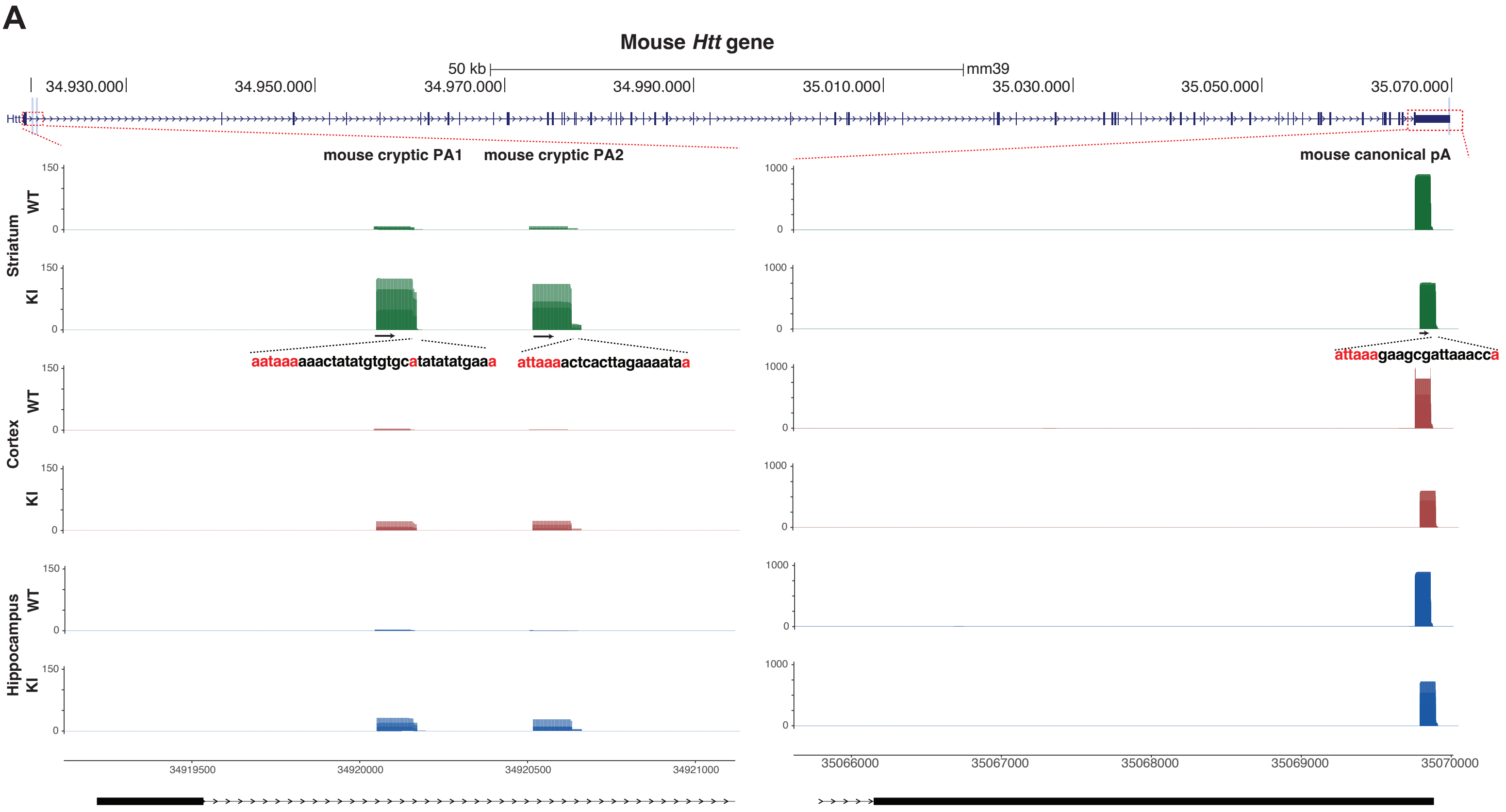

### Supplementary Figure 4

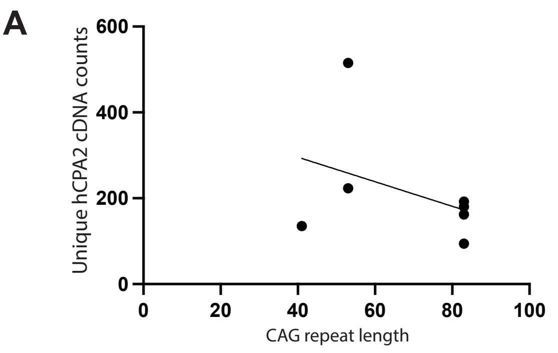


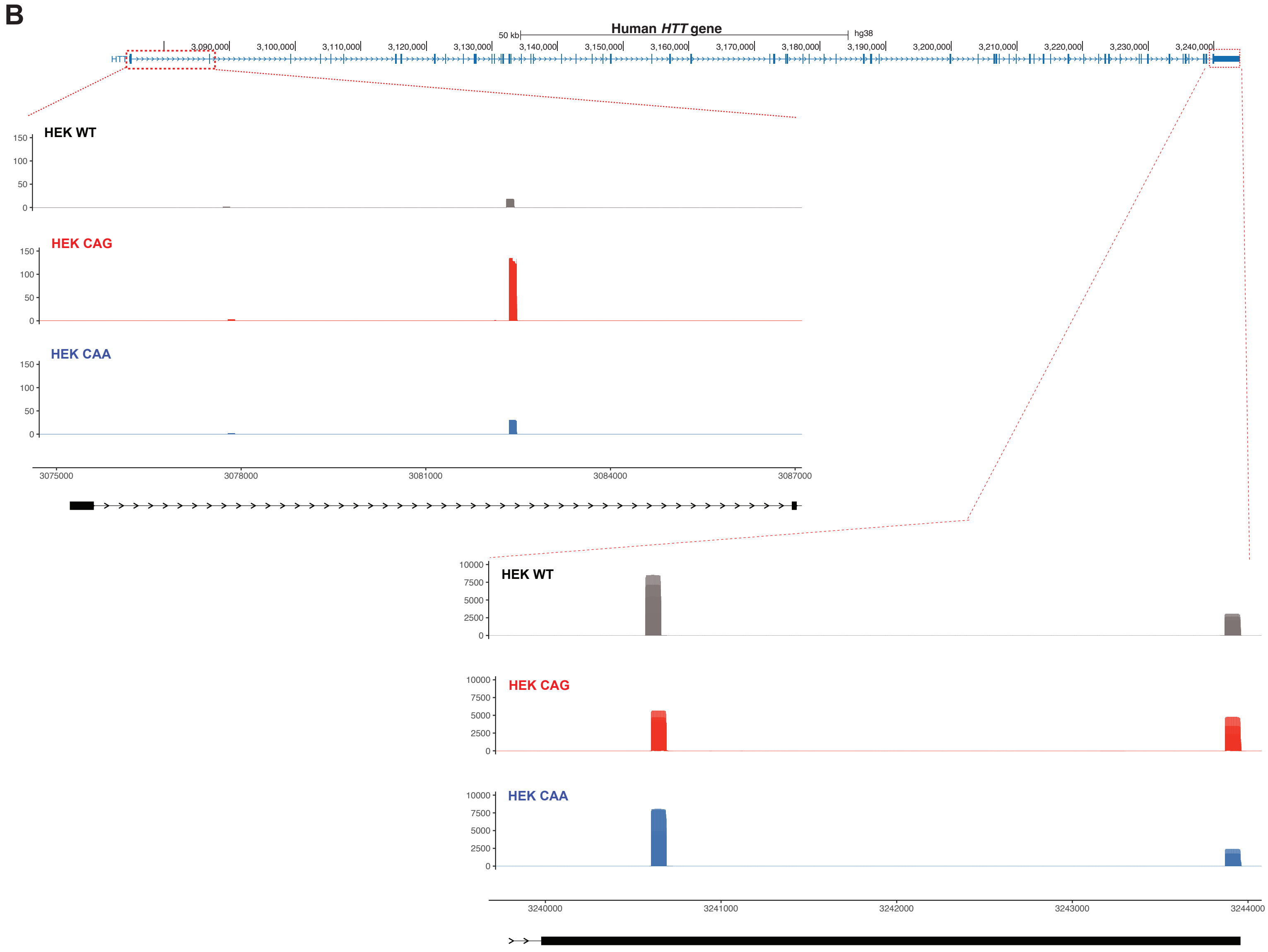

### Supplementary Figure 5

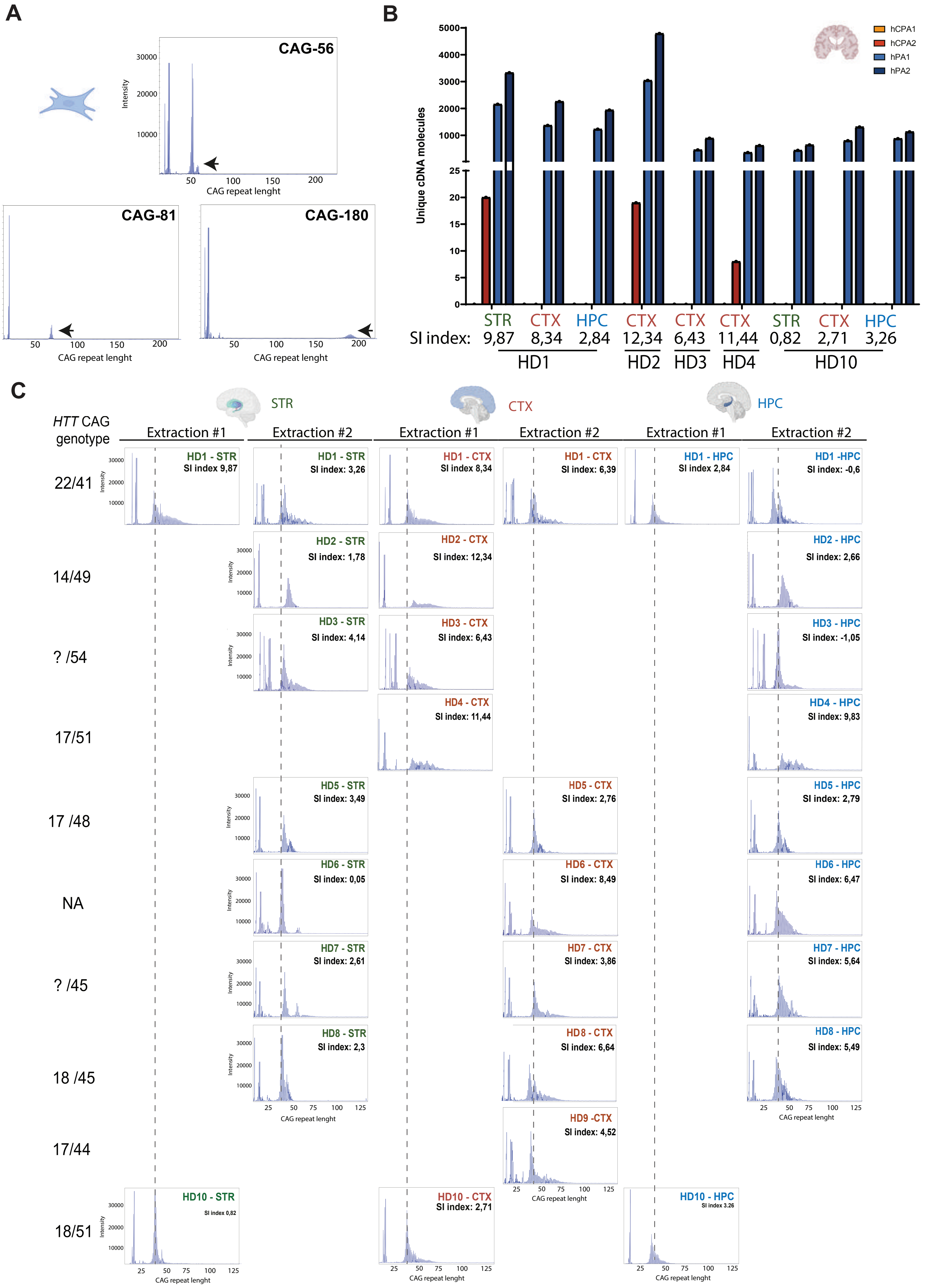
