## Supplementary Material for "Targeted 3’-end RNA sequencing uncovers cryptic polyadenylation in Huntington’s disease linked to somatic instability and CAG repeat purity"

**Supplementary figure legends**

**Supplementary Figure 1. Related to Figure 1. A.** Sashimi plots of WT and YAC128 mouse brain regions showing the reads mapping the expected *HTT* canonical 3´-ends. The arrow indicates the location of the oligonucleotide used for the second strand synthesis. The sequence at the end of the read indicates the polyadenylation signal (PAS) and the cleavage sites in red. **B.** Unique cDNA read counts for human *HTT* 3´-ends indicated in the y-axis of each plot after performing second strand synthesis with the oligonucleotides shown in the x-axis. **C.** Unique cDNA read counts of human *HTT* and mouse *Htt* transcripts in the three different brain regions analyzed in the YAC128 HD mouse model. Data are presented as mean ± SD. Statistical significance was calculated with a one-way ANOVA test followed by Tukey´s multiple comparisons: ns (non-significant), * (P ≤ 0.05), ** (P ≤ 0.01) or *** (P ≤ 0.001).

**Supplementary Figure 2. Related to Figure 2. A.** Unique cDNA read counts of cryptic and canonical human *HTT* and mouse *Htt* transcripts obtained in singleplexed versus multiplexed reactions in the three different brain regions analyzed in the YAC128 HD mouse model. Data are presented as mean ± SD. Statistical significance was calculated using an unpaired Student´s t-test: ns (non-significant). **B.** Sashimi plots of YAC128 mouse brain regions showing the reads mapping the expected *HTT* 3´-ends.

**Supplementary Figure 3. Related to Figure 3. A.** Sashimi plots of Hdh^Q111^**^/+^** mouse brain regions showing the reads mapping the expected *Htt* 3´-ends. The arrow indicates the location of the oligonucleotide used for the second strand synthesis. The sequence at the end of the read indicates the polyadenylation signal (PAS) and the cleavage sites in red.

**Supplementary Figure 4. Related to Figure 4. A.** Simple linear regression analysis of the number of CAG repeats in HEK293-CAG cell lines (N=7) versus unique cDNA read counts for the distal cryptic *HTT* transcript. **B.** Sashimi plots of the three different HEK293 cell lines used in this study, mapping the expected *HTT* 3´-ends.

**Supplementary Figure 5. Related to Figure 5. A.** Length distribution of *HTT* CAG tract in three human HD fibroblasts. **B.** Unique cDNA read counts of cryptic and canonical *HTT* transcripts in 9 human HD brain regions analyzed by 3TRS starting with 1μg RNA. Data are represented as individual values. **C.** Length distribution of *HTT* CAG tract across the RNA samples isolated from HD postmortem brain tissues used in this study. The vertical dashed line sets the 40 CAG repeat length threshold. Extraction #1 corresponds to the RNA samples used in the 3TRS experiment performed with 1ug RNA and shown in Fig. Sup 5B while Extraction #2 correspond to the additional RNA samples isolated afterwards to perform 3TRS with a 4ug RNA input and shown in Fig. 5D. Each fragment analysis includes the associated SI index. The *HTT* CAG repeat length for each HD case, determined from the genomic DNA isolated from peripheral blood at the time of genetic diagnosis, is indicated in the left column.

**Supplementary Tables**

**Supplementary Table 1.** Human postmortem brain samples used in this study. Cortex (CTX), hippocampus (HPC) and striatum (STR). NA= data not available.

| Code | Brain regions | Sex | Age at death | CAG number at diagnosis |
| --- | --- | --- | --- | --- |
| HD1 | CTX, HPC, STR | Male | 61 | 22/41 |
| HD2 | CTX, HPC, STR | Male | 43 | 14/49 |
| HD3 | CTX, HPC, STR | Male | 34 | NA/54 |
| HD4 | CTX, HPC | Male | 47 | 17/51 |
| HD5 | CTX, HPC, STR | Male | 57 | 17/48 |
| HD6 | CTX, HPC, STR | Male | 48 | NA |
| HD7 | CTX, HPC, STR | Male | 53 | NA/45 |
| HD8 | CTX, HPC, STR | Female | 56 | 18/45 |
| HD9 | CTX | Male | 58 | 17/44 |
| HD10 | CTX, HPC, STR | Female | 62 | 18/51 |

**Supplementary Table 2.** Engineered HEK293 HD cell lines used in this study.

| Short Name | *HTT* CAG Genotype |
| --- | --- |
| Hek293-WT-1 | 16/17 |
| Hek293-WT-2 | NA |
| Hek293-WT-3 | NA |
| Hek293-WT-4 | 0/0 |
| Hek293-HD-1 | 41/41 |
| Hek293-HD-2 | 53/53 |
| Hek293-HD-3 | 53/53 |
| Hek293-HD-4 | 83/83 |
| Hek293-HD-5 | 83/83 |
| Hek293-HD-6 | 83/83 |
| Hek293-HD-7 | 83/83 |
| Hek293-CAA-1 | 40CAG plus 1xCAA/40CAG plus 1xCAA |
| Hek293-CAA-2 | 39CAG plus 2xCAA/39CAG plus 2xCAA |
| Hek293-CAA-3 | 39CAG plus 2xCAA/39CAG plus 2xCAA |
| Hek293-CAA-4 | 39CAG plus 2xCAA/39CAG plus 2xCAA |
| Hek293-CAA-5 | 39CAG plus 2xCAA/39CAG plus 2xCAA |

**Supplementary Table 3.** Oligonucleotides used in this study.

| Oligo name | Sequence 5´-3´ | Use |
| --- | --- | --- |
| RT_oligo_dT | 5´ GTTCAGACGTGTGCTCTTCCGATCT**NNNNN**(T)20 3´ | For RNA retrotranscription. The **N** nucleotide indicate the barcode sequence that allows sample multiplexing after RT |
| *HTT*_PA1 | 5´ CACGACGCTCTTCCGATCT**N**_(6/10)_gctagacacccggcaccatt 3´ | For second strand synthesis. The common sequence in upper case contains Illumina adaptors.  The **N** nucleotide represents the random nucleotides of the unique molecule identifier (UMI). The nucleotides. The sequence is lower case is specific for each interrogated 3´ end. |
| *HTT*_PA2 | 5´ CACGACGCTCTTCCGATCT**N**_(6/10)_gcaaagggaaggactgacgaga 3´ |  |
| *HTT*_cPA1 | 5´ CACGACGCTCTTCCGATCT**N**_(6/10)_gccaccatgcctggctagaat 3´ |  |
| *HTT*_cPA2 | 5´ CACGACGCTCTTCCGATCT **N**_(6/10)_ccccatcacctccatttcctgt 3´ |  |
| *Htt*_PA | 5´ CACGACGCTCTTCCGATCT**NNNNNN**tagacctggctgtccatttccaaaa 3´ |  |
| *Htt*_cPA1 | 5´ CACGACGCTCTTCCGATCT**NNNNNN**ccagagaggtgtgcttctgtgt 3´ |  |
| *Htt*_cPA2 | 5´ CACGACGCTCTTCCGATCT**NNNNNN**cctttctctcaactctgggc 3´ |  |
| P5-Fwd | 5´ AATGATACGGCGACCACCGAGATCTACACTCTTTCCCTAC  ACGACGCTCTTCCGATCT 3´ | Oligos for final PCR reaction. The **N** nucleotide represents the i7 index for Illumina demultiplexing |
| P7-Rv | 5´ CAAGCAGAAGACGGCATACGAGAT**NNNNNN**GTGACTGG  AGTTCAGACGTGTGCTCTTCC 3´ |  |
